# Dynamic Changes in Macrophage Polarization during the Resolution Phase of Periodontal Disease

**DOI:** 10.1101/2023.02.20.529313

**Authors:** Juhi R. Uttamani, Varun Kulkarni, Araceli Valverde, Raza Ali Naqvi, Salvador Nares, Afsar R. Naqvi

## Abstract

Periodontal inflammation is largely governed by infiltration of myeloid cells, in particular macrophages. Polarization of Mφ within the gingival tissues is a well-controlled axis and has considerable consequences for how Mφ participate in inflammatory and resolution (tissue repair) phases. We hypothesize that periodontal therapy may instigate a pro-resolution environment favoring M2 Mφ polarization and contribute towards resolution of inflammation post-therapy. We aimed to evaluate the markers of macrophage polarization before and after periodontal therapy. Gingival biopsies were excised from human subjects with generalized severe periodontitis, undergoing routine non-surgical therapy. A second set of biopsies were excised after 4-6 weeks to assess the impact of therapeutic resolution at the molecular level. As controls, gingival biopsies were excised from periodontally healthy subjects, undergoing crown lengthening. Total RNA was isolated from gingival biopsies to evaluate pro- and anti-inflammatory markers associated with macrophage polarization by RT-qPCR. Mean periodontal probing depths, CAL and BOP reduced significantly after therapy and corroborated with the reduced levels of periopathic bacterial transcripts after therapy. Compared to heathy and treated biopsies, higher load of Aa and Pg transcripts were observed in disease. Lower expression of M1Mφ markers (TNF-α, STAT1) were observed after therapy as compared to diseased samples. Conversely, M2Mφ markers (STAT6, IL-10) were highly expressed in post-therapy as opposed to pre-therapy, which correlated with clinical improvement. These findings corroborated with murine ligature-induced periodontitis and resolution model, comparing the respective murine Mφ polarization markers (M1 Mφ: *cox2*, *iNOS2* and M2 Mφ: *tgm2* and *arg1*). Our findings suggest that imbalance in M1 and M2 polarized macrophages by assessment of their markers can provide relevant clinical information on the successful response of periodontal therapy and can be used to target non-responders with exaggerated immune responses.

## Introduction

The classic chronic inflammatory state in periodontal disease is characterized by prolonged inflammation in conjunction with healing events *viz*. angiogenesis and fibrosis ^1^. The inflammatory microenvironment in periodontal disease is governed by infiltration of myeloid cells along-with the local stromal macrophages (MΦ) help to augment immune cell activation and prevent tissue damage. MΦ can exhibit inherent functional plasticity via initiating the innate responses and shaping the adaptive arm of immunity allowing them to thus, adeptly respond according to a myriad of pathogenic stimulus in the oral cavity. The phagocytic abilities of both tissue-resident MΦs and monocyte-derived recruited MΦ alike, are essential in innate defense as well as in the regulation of acquired immunity. Studies report differential transcriptional profiling in the tissue resident versus recruited MΦ from circulating monocytes, signifying that location and environmental triggers stimulate their polarization ^2^. Based on the type-1/type-2 T helper cell polarization concept, phenotypically polarized MΦ can differentiate into M1 (classical) or M2 (alternative) phenotypes ^3^. Exemplifying opposing activities, M1 MΦ promote bacterial killing and a proinflammatory environment by increased production of IL-6, TNF-α, inducible nitric oxide synthase (iNOS), etc., while M2 MΦ are anti-inflammatory in nature favoring resolution of inflammation and wound-healing.

Preliminary studies from our lab have shown that the priming of naive monocytes to host cytokines, such as interferon gamma (IFN-γ) or IL-4, can give rise to different MΦ phenotypes ^4^. This has considerable consequences for how MΦ detect, phagocytose and kill bacteria. MΦ polarization in pathologic conditions such as chronic periodontitis most appropriately represents a continuum, as opposed to two distinct scenarios involving heterogeneity of the effector cells to a spectrum of polarization states that do not fit to the oversimplified M1/M2 classification ^5^. The local microenvironment dependent abundance of MΦ within tissues is finely controlled through the axis of M-CSF and GM-CSF, enabling appropriate responses to the microbial challenge or repair following an injury.

The inflammatory microenvironment in periodontal disease is governed by infiltration of myeloid cells along-with the local stromal MΦ help to augment immune cell activation and prevent tissue damage. Periopathogens (such as *P. gingivalis* and *A. actinomycetemcomitans*) alter host immune response by perturbing expression of pro-and anti-inflammatory milieu causing activation of M1MΦ phenotype that are primarily involved in Th1 responses, lymphokine production, and degradation of intracellular pathogens. Periodontal therapy that results in reduction of bacterial load that may instigate a pro-resolution environment favoring M2 MΦ polarization, triggering Th2 responses such as immunotolerance, and tissue remodeling, etc.

Human MΦ respond to live Pg or Aa and their LPS by polarizing towards the pro-inflammatory M1 phenotype ^4^. We hypothesized that non-surgical periodontal therapy creates a M2MΦ dominant microenvironment favoring tissue repair. In this study, the expression of bacterial gene transcripts from Pg and Aa that were most consistently expressed in gingival biopsies of chronic periodontitis subjects across various geographically distinct cohorts (*Naqvi et al., Unpublished results*) were examined to reflect their changes post-therapy. There have been reports studying the expression of M1, M2 markers in human gingival tissues of patients with chronic periodontitis ^6^. However, to the best of our knowledge there has been no longitudinal study evaluating the influence of periodontal therapy on MΦ polarization. Our aim was to fill existing knowledge gaps by examining the profiles of polarized MΦ (M1 and M2) after non-surgical periodontal therapy to provide further insights of their role in modulating oral mucosal immune responses as well their active participation in wound repair. This transition between M1 and M2 polarization can ultimately determine the pathogenesis of periodontal disease, given the dynamic interaction of immune mediated signaling governing the inflammatory component as well as tissue healing in the cyclical nature of periodontal disease. We anticipate that these changes can provide insightful findings on the role pathogen stimulated immune responses in the development of oral diseases. The ultimate goal of which being identification of therapeutically beneficial drug targets that will enhance current modalities for the treatment of periodontitis.

## Materials and Methods

### Subject selection and sample collection

The present investigation was an observational, cross-sectional pilot study approved by the Institutional Review Board and the Ethics Research Committee at the University of Illinois at Chicago, College of Dentistry (IRB Protocol# 2017-1064). The study was conducted according to the ethical principles of the Helsinki Declaration. Inclusion criteria included male and female patients ages 18–65 years and in good systemic health. Exclusion criteria included chronic disease (diabetes, hepatitis, renal failure, clotting disorders, HIV, etc.), smokers, antibiotic therapy for any medical or dental condition within a month before screening, and subjects taking medications known to affect periodontal status (e.g., phenytoin, calcium channel blockers, cyclosporine). Briefly, subjects presenting to the Postgraduate Periodontics Clinic at the UIC College of Dentistry for routine periodontal care from March 2018 to May 2019 were recruited for this study. All participants were informed of the aims of the study and signed the informed written consent form prior to entering the study. The convenience sample from the present study comprised twelve individuals from a multiethnic group, of both sexes, and divided into two groups: periodontally healthy (Control, H; n = 12) or chronic periodontitis (Experimental, D; n = 12).

For the diseased group, a biological sample (gingival tissue including the gingival epithelium, col area and underlying connective tissue) were obtained at baseline (D, pre-therapy) and 4-6 weeks after non-surgical periodontal therapy (T, post-therapy) from the same subject. The second biopsy was taken from a separate, preselected site. For the control group (H), samples were derived from healthy gingival tissues normally discarded during routine crown lengthening procedures. Periodontal status was assessed using periodontal probing depth (PD), clinical attachment level (CAL), bleeding on probing (BOP), and radiographic evaluation of crestal bone loss. All examinations were performed with a manual periodontal probe (PCPUNC-15, Hu-Friedy, Chicago, IL, USA) by two investigators (J.R.U. and V.K), with high inter-examiner correlation coefficient kappa ‘0.85’. The presence or absence of chronic periodontitis was described by our previous studies ^8^. Briefly, subjects with generalized severe chronic periodontal disease displayed probing depth ≥ 6 mm, ≥ 4mm of attached gingiva, bleeding on probing and radiographic evidence of bone loss ^9^. Health periodontal patients displayed probing depths ≤ 3 mm, with no bleeding on probing and no radiographic evidence of bone loss ^10^.

### Total RNA Isolation and RT-qPCR

Tissue samples were lysed using the TissueLyzer (Qiagen) and total RNA isolated using the miRNeasy kit (Qiagen), with the subsequent mRNA and miRNA expression analysis as previously reported ^8,11^. Expression levels of STAT1, STAT6, TGM2, CXCL10, CCL22, IL10 and normalization control GAPDH were examined by RT-qPCR using gene specific primers (Sigma Aldrich, St. Louis, MO, USA) as previously described ^11^. Real-time PCR based analysis was also used to assess the impact of therapy on the load of periodontal pathogens: *Actinobacillus actinomycetemcomitans, Porphyromonas gingivalis* by testing bacterial gene products of the two most highly expressed candidate genes for each. Briefly, transcript levels of primers designed for gingipain RgpA, RNA polymerase sub-unit β of *P. gingivalis* and Elongation Factor Tu, Ribosomal protein L2 of *A. actinomycetemcomitans* were evaluated by RT-PCR (***Table 1***).

**Table 1.**
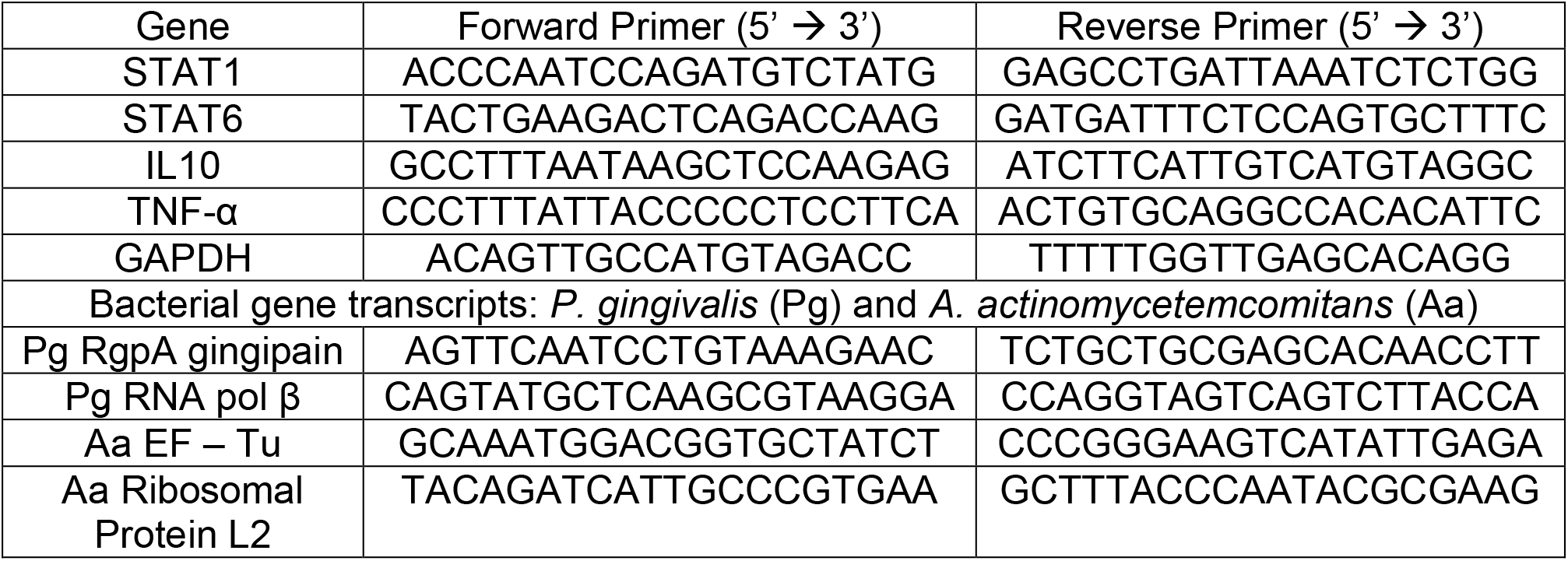
Primer sets used for M1, M2 macrophage markers and bacterial gene transcripts.

Additionally, for mature miRNA-155 quantification, miScript primers and miScript II RT Kit were purchased from Qiagen. 100 ng total RNA was reverse transcribed according to manufacturer’s instruction. The reactions were performed using miRNA specific primer, universal primer (Qiagen) and SYBR Green (Roche, Indianapolis, IN, USA). RNU6B was used as endogenous control. The Ct values of replicates were analyzed to calculate relative fold change using the delta-delta Ct method and the normalized values plotted as histograms with standard deviations (SD). The Ct values of replicates were analyzed to calculate relative fold change using the delta-delta Ct method. The primers used for our experiments are listed in ***Table 1***.

### Murine model of ligature-induced periodontitis and its resolution

Periodontitis was induced using a 6-0 silk ligature placed bilaterally between the maxillary first and second molar in 8-12-week-old female mice (n=4/group) under anesthesia using intraperitoneal injection of ketamine and xylazine. Animals were euthanized at 8 day post-ligature (DPL) placement. For resolution of periodontal inflammation, we removed the ligature on day 8 and sacrificed the animals on day 10 post-ligature removal. The mice with displaced ligature during experimental period were excluded from the evaluation. Gingival tissue were harvested from around the maxillary first and second molars and homogenized. Total RNA was extracted from the excised gingival tissue using the miRNeasy kit (Qiagen). RT-qPCR was performed as described above and the gene expression levels were normalized to those of the reference endogenous controls β-actin.

### Statistical Analysis

Data were analyzed using GraphPad Prism (GraphPad Software, La Jolla, CA, USA). The results were presented as SD or ±SEM from three independent replicates, and experiments were conducted at least three times. The p values were calculated using a Student *t* test and/or ANOVA for more than two groups. A p value <0.05 was considered significant. Age distribution in the healthy controls and disease groups were assessed by Mann-Whitney U test. Gender distribution across the groups was evaluated by McNemar χ2 test.

## RESULTS

### Non-surgical periodontal treatment improves clinical parameters

The sample size of this pilot study consisted of 12 individuals. Accounting for the pre-treatment and post-treated samples the total number of 24 biologic replicates for the three groups were as follows: Healthy Controls ‘H’ n = 12, Diseased ‘D’ n = 12, Treated ‘T’ n = 12, respectively. Periodontal clinical parameters of study groups and the subject demographics are shown in ***Figure 1A***. There were no statistically significant differences noted in age (p=0.49) or gender (p=0.55) across the groups in our cohort. The mean age of individuals in the healthy group being 43 ± 11.3 years (range: 32–60) while the mean age in the chronic periodontitis/treated group was 52 ± 9.31 years (range: 38–59). A statistically significant reduction in the Periodontal Pocket Depth (PPD), Clinical Attachment loss (CAL) and Bleeding of Probing (BOP) was observed post-therapy (***Figure 1 B-D***). For individuals with generalized chronic severe periodontitis, mean pocket depth (PD) was 9 mm pre-therapy, which was reduced to approximately 7 mm after the non-surgical periodontal therapy (***Figure 1B***). The mean PPD in the control group was 2.75 mm. We also noticed significant reduction in mean CAL (5.59 ± 0.35 mm; p<0.01) and %BOP (25.0 ± 7.35) in post-therapy compared to pre-therapy (9.35 ± 1.28 mm) (***Figure 1 B,C***). For periodontally healthy subjects, CAL and BOP values were 3.15 ± 0.12 mm and 7 ± 2.4. Overall, non-surgical periodontal therapy improves clinical parameters in our cohort.

**Figure 1.**
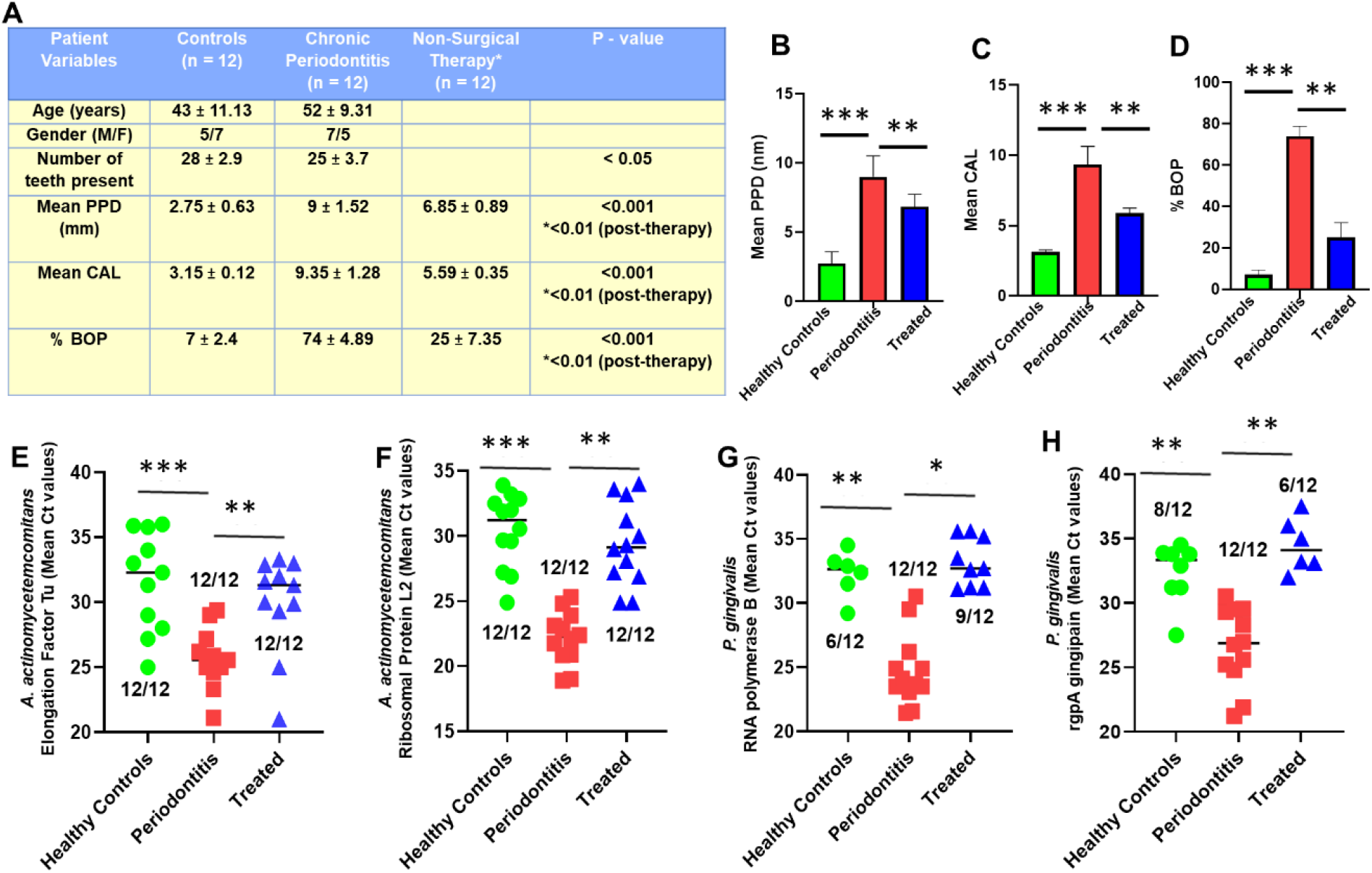
Non-surgical periodontal therapy improves clinical parameters by reducing gingival periopathogen burden. (A) Subject demographics and clinical parameters: The total sample size of the study was 24 samples included from the 3 groups: Healthy N = 12, Diseased N = 12 and Treated N = 12. There were no statistically significant differences in age and gender between the groups. Reduction in the mean (B) periodontal probing depth, (C) clinical attachment loss and (D) percent bleeding on probing (BOP) were observed in the diseased individuals as measured 4-6 weeks after non-surgical periodontal therapy (p<0.05). Mann-Whitney U test was used to compare age and McNemar’s Chi-square test was used to compare gender distribution between the two groups. Age is presented as mean ± standard deviation. Reduced levels of Aa- and Pg-encoded gene transcripts post-therapy. Quantitative RT-PCR showing mean Ct values of Aa-encoded (E) Elongation Factor-Tu and (F) Ribosomal protein L2 and Pg-encoded (G) RNA polymerase subunit-b and (H) RgpA gingipain in gingival biopsies after periodontal therapy. Numbers of samples showing detection of transcripts (cut-off value Ct>37) are showing for each group. Student’s t-test was used to calculate p-values. *p < 0.05, **p < 0.01, ***p < 0.001. Data is presented as ±SEM of all the detectable readings.

### Reduced levels of periodontopathic bacterial transcripts in gingival biopsies after periodontal therapy corroborates with improvement in clinical parameters

The goal of non-surgical periodontal therapy encompasses removal of plaque biofilm and calculus via scaling and root planning. To investigate the impact of non-surgical periodontal therapy response at the molecular level on the bacterial load, we compared expression of bacterial gene transcripts in gingival biopsies derived from periodontally healthy, diseased, and treated subjects. Our preliminary analysis indicated RgpA gingipain (PG 2024), RNA poly β (rpoB) (PG 0395) to be one of the most highly expressed Porphyromonas gingivalis encoded transcripts enriched in chronic periodontitis. Likewise, Elongation Factor – Tu (EF – Tu)/ AA 918 and Ribosomal protein L2 (RPL2)/ AA 778 were two of the most significantly enriched transcripts encoded by Aggregatibacter actinomycetemcomitans (Aa), as observed by RNA sequencing analysis (RNAseq) (*Naqvi et al., Unpublished data*). This observation was consistent across two different, geographically distinct cohorts of chronic periodontitis subjects.

Hence, we attempted to examine the expression levels of RgpA, rpoB, EF-Tu and RPL2 in the gingival biopsies that were collected from subjects diagnosed with chronic periodontitis and compare it after non-surgical periodontal therapy (4-6 weeks post-treatment) and from healthy controls. Our data confirmed significantly higher expression of *A. actinomycetemcomitans* transcripts EF-Tu (mean Ct = 25.66 ± 2.37), RPL2 (mean Ct = 22.15 ± 2.03) (***Figure 1E,F***) and *P. gingivalis* transcripts rpoB (mean Ct = 24.74 ± 2.8), RgpA (mean Ct = 26.63 ± 3.01) (***Figure 1G,H***) in diseased biopsy specimens as compared to healthy controls or post-treatment samples. Both the *A. actinomycetemcomitans* and *P. gingivalis* gene transcripts were expressed at low levels or could not be detected in some samples in the healthy gingival biopsies. For the gingival biopsies taken after 6 weeks post non-surgical therapy, the levels of the bacterial gene transcripts approximated as that seen in healthy controls, for both the periodontal pathogens. In post-therapy samples, Aa transcripts EF-Tu (mean Ct = 29.93 ± 3.78) and RPL2 (mean Ct = 29.35 ± 3.16) were detected at low levels. Pg transcripts rpoB (mean Ct = 33.19 ± 1.88) and RgpA (mean Ct = 34.47 ± 2.06), were either not detected or showed significantly reduced expression levels post-therapy. These results clearly show that the transcripts of periodontal pathogens demonstrated significantly greater expression in gingival biopsies derived from periodontally diseased subjects and its expression was reduced after periodontal therapy, supporting a central role of bacterial pathogens in periodontal pathology.

### Downregulation of M1 macrophage marker expression after non-surgical periodontal therapy

To investigate whether non-surgical periodontal therapy can impact MΦ polarization we evaluated the changes in expression of well M1 markers STAT1 and miR-155, that have been well characterized to promote M1 MΦ phenotype ^12,13^. We compared expression of STAT1 and miR-155 in gingival biopsies derived from periodontally healthy, diseased, and treated (4-6 weeks post-treatment) subjects. We observed significantly higher expression of STAT1 (Fold change ~4) and miR-155 (Fold change ~3) in diseased gingival samples compared to healthy controls (***Figure 2 A,B***). The expression levels of STAT1 and miR-155 in gingival biopsies taken after 6-weeks post non-surgical therapy were significantly decreased, similar to levels observed in the healthy cohort (***Figure 2***).

Likewise, upon assessing M1 related pro-inflammatory cytokine/chemokine expression, we found higher expression of TNFα (Fold change ~ 4) and CXCL10 (Fold change ~3.5) in samples from chronic periodontitis subjects as compared to healthy controls (***Figure 2C,D***). The expression of both TNFα and CXCL10 significantly reduced in post-treatment biopsies. These results clearly show that M1 MΦ markers are downregulated in gingival biopsies derived after non-surgical periodontal therapy from periodontally diseased subjects.

**Figure 2.**
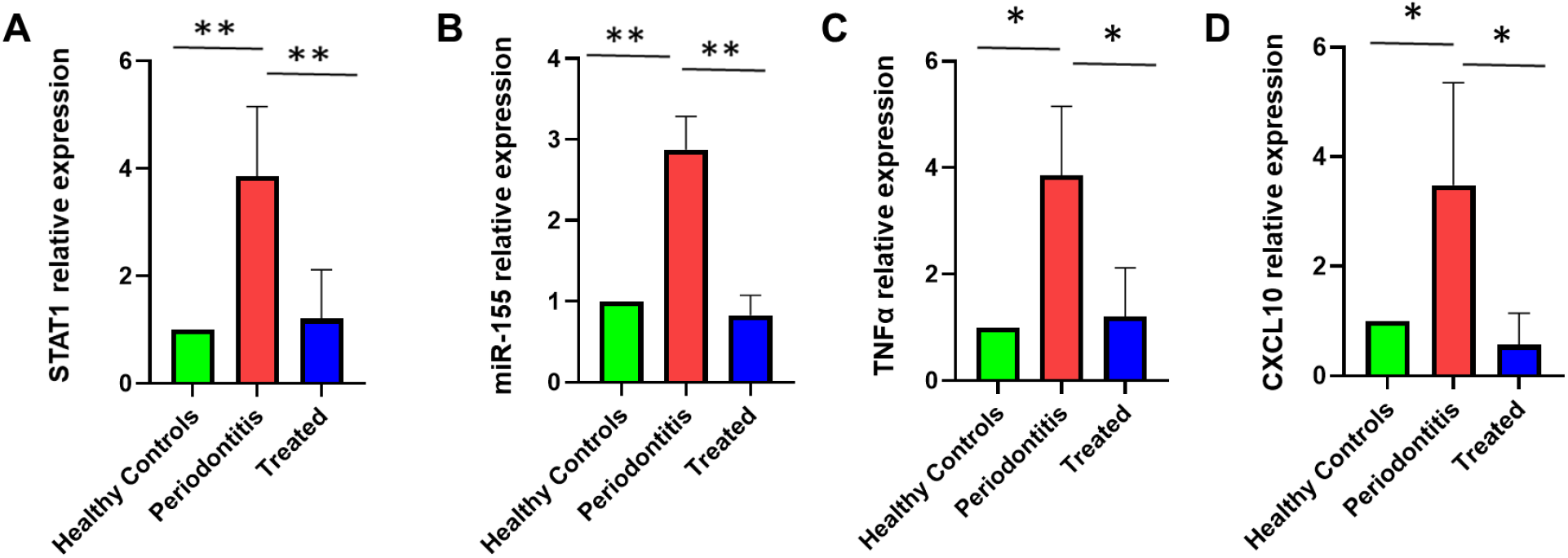
Downregulation of M1 macrophage markers after non-surgical therapy. Histograms showing relative expression of pro-inflammatory (A) STAT1, (B) miR-155, (C) TNF-α and (D) CXCL10 transcripts by quantitative PCR analysis in gingival biopsy samples of periodontitis patients (n = 12), after therapy (n = 12) as compared to those of healthy controls (n = 12). Results are normalized to those of controls and are represented relative to expression of GAPDH and RNU6, respectively. Student’s t-test was used to calculate p-values. *p < 0.05, **p < 0.01. Data is presented as ±SEM of all samples.

### Upregulation of gingival M2 macrophage markers after non-surgical periodontal therapy

To theorize the antagonistic expression levels of M2MΦ activation markers in response to periodontal therapy, we evaluated expression profiles of TGM2 and STAT6, which have been well characterized to promote M2 phenotype ^12,14^. We also examined the expression of M2 associated effector chemokines/cytokines (CCL22, IL10) in response to therapy. We inspected the RNA expression levels of STAT1, TGM2, IL10, and CCL22 in the gingival biopsy specimens from both the periodontally diseased cohorts as well as samples obtained post-treatment. In contrast to the elevated expression of M1 markers as previously observed, we found diminished expression of M2 markers STAT6 (Fold change ~10) and TGM2 (Fold change ~8) in the diseased specimens as compared to healthy controls as verified by quantitative PCR (***Figure 3 A,B***). Conversely, periodontal therapy recuperated STAT6 (Fold change ~4; compared to disease) and TGM2 (Fold change ~3.5; compared to disease) expression levels and their expression were significantly higher compared to pre-therapy group. Likewise, upon assessing M2 associated anti-inflammatory cytokine/chemokine expression, we found higher expression of IL10 (Fold change ~2) and CCL22 (Fold change ~6) in post-treatment samples (***Figure 3C,D***). Expression of anti-inflammatory cytokines IL10 (Fold change ~8), and CCL22 (Fold change ~8) were significantly less in the pre-treatment diseased biopsies. Collectively, these demonstrate that M2MΦ or the repair-associated markers are upregulated in gingival biopsies harvested after non-surgical periodontal therapy from periodontally diseased subjects.

**Figure 3.**
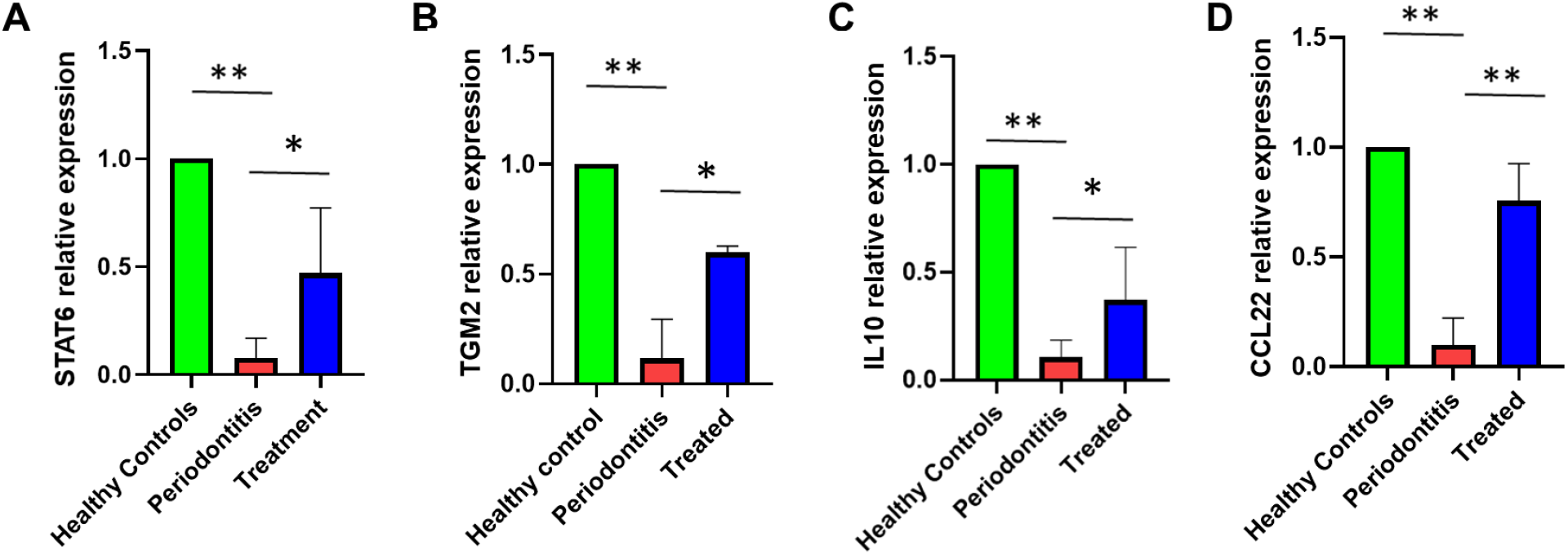
M2 macrophage markers are upregulated after non-surgical therapy. Histograms showing relative expression of anti-inflammatory markers (A) STAT6, (B) TGM2, (C) IL10 and (D) CCL22 by quantitative RT-PCR analysis in gingival biopsy samples of periodontitis patients (n = 12), after therapy (n = 12) as compared to those of healthy controls (n = 12). Results are normalized to those of controls and are represented relative to expression of GAPDH. Student’s t-test was used to calculate p-values. *p < 0.05, **p < 0.01. Data is presented as ±SEM of n=12 /group.

### Macrophages exhibit dynamic changes in the expression of phenotype markers in a murine model of ligature-induced periodontitis and resolution

Our human studies suggest that M1 and M2 phenotype exhibit dynamic changes pre- and post-therapy. To confirm these findings, we examined M1 and M2 markers expression in gingival biopsies collected from mice subjected to ligature-induced periodontitis. Placement of ligature alone can cause dysbiosis and induce periodontal inflammation and marked alveolar bone loss. For resolution, we removed the ligature on day 8 (disease established) in a parallel animal cohort to activate resolution process for 10 day (DPLR [day post-ligature removal]. ***Figure 4A*** shows schematic that describes the inflammation and resolution murine model. We observed significantly higher murine M1 markers *cox2* (Fold change ~6) and *inos2* (Fold change ~8) in the gingiva at 8DPL, which marks disease establishment (***Figure 4B,C***). Interestingly, after 10 day post-ligature removal (DPLR) we noticed significantly less expression of inflammatory markers *cox2* (Fold change ~6 vs 8DPL) and *inos2* (Fold change ~8 vs 8DPL) compared to 8DPL and shows similar levels observed for no ligature control. These results suggest a reversal in macrophage phenotype after removal of inflammatory stimuli.

**Figure 4.**
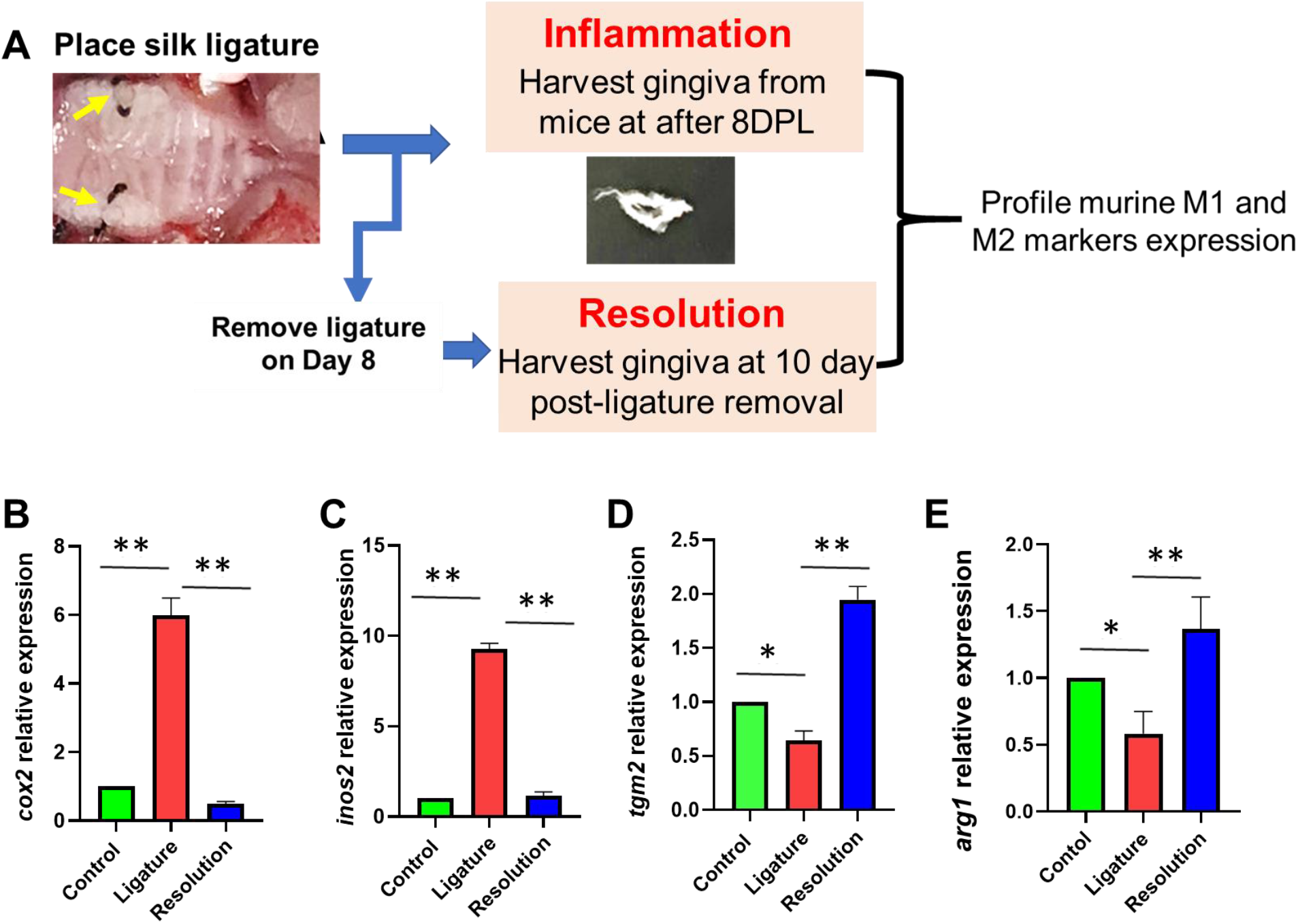
M1 and M2 macrophage markers exhibit antagonistic expression in murine ligature-induced periodontitis and resolution model. (A) Schematic showing murine periodontal inflammation and its resolution model. Total RNA was isolated from healthy and inflamed gingival biopsies from mice (n=4/group) subjected to ligature-induced periodontitis for 8 days and after 10 day post-ligature removal. Expression of M1 and M2 markers was quantified by RT-qPCR. Histograms showing relative fold change expression of (B) *cox-2*, (C) *iNOS2*, (D) *tgm2* and (E) *arg1* in murine gingiva. Transcript expression was normalized to β-actin. Student’s t-test was used to calculate p-values. *p < 0.05, **p < 0.01. Data is presented as ±SEM of n=4/group.

We next asked whether M2 markers show opposite pattern in our murine model as observed in inflamed and post-therapy human gingiva. Our results show significantly reduced levels transglutaminase 2 (*tgm2;* Fold change ~0.75) and arginase1 (*arg1;* Fold change ~0.5) compared to animals without ligature (***Figure 4D,E***). On the contrary, 10 DPLR group showed marked increase in both *tgm2* (fold change ~2.5) and *arg1* (fold change ~2) levels compared to 8DPL indicating a pronounced activation of pro-resolution M2 phenotype. Overall, these results strongly support that M1 and M2 phenotype markers in gingiva exhibit antagonism after inflammation and during resolution of inflammation *in vivo* indicating a functional role of macrophage polarization in periodontal tissue immune homeostasis.

## Discussion

Chronic periodontitis is a polymicrobial, inflammatory disease that commences with periodontal pathogen-induced immune responses, affecting the supporting tissues of the teeth including the periodontium and the alveolar bone ^15,16^. Based on the concept of immuno-modulation, macrophages play important roles as effector cells in mediating Th1 driven and Th2 derived immune responses. It is not incorrect to state that the chronicity of periodontal inflammation is an aggregated set of events with a continued proinflammatory environment accompanied with cyclic bursts of tissue healing patterns, including cell apoptosis, wound healing, cell proliferation, tissue repair ^16^. Upcoming literature evidence suggests the profound role of macrophage polarization in the gingival tissue from chronic periodontitis subjects ^5,13,17^. However, to the best of our knowledge there has been no study undertaken to longitudinally characterize MΦ polarization after non-surgical periodontal therapy in humans.

To address this shortcoming, we designed a case-control pilot study using a representative cohort of 24 subjects ranging from 19 to 80 years old presenting with no clinical history of uncontrolled systemic disease or use of drugs that could have an impact the periodontal disease pathogenesis or influence therapy. We also excluded pregnant subjects as well as patients with a history of uncontrolled diabetes, use of antibiotics, and smokers. To explore a more accurate picture from the standpoint of periodontal pathogenesis and minimize potential confounding factors (e.g. age, gender), we focused on a cohort of patients presenting chronic generalized severe periodontal disease with age and gender matched controls. We evaluated the global macrophage polarization through assessing changes observed between M1 and M2 MΦ markers in the gingival tissue samples of periodontally diseased subjects before and after therapy and compared it with healthy controls. In agreement with previous studies, the treatment of subjects in the chronic periodontitis cohort by scaling and root planning in promoting a clinically favorable environment, resulted in significant improvements in probing depths, clinical attachment levels and bleeding on probing parameters ^18,19^. In additional, to compare the findings from human gingival biopsies and add robustness to our data, we also designed a murine model of periodontal disease and resolution.

Studies have reported higher prevalence of *P. gingivalis* and *A. actinomycetemcomitans* in subjects with chronic periodontitis ^20^. Longitudinal studies assessing the clinical and microbiological evaluation of individuals undergoing periodontal therapy, have observed reduction in bacterial loads in the subgingival sites post-therapy ^21,22^. However, both Pg and Aa being able to invade the host tissues (e.g.gingival epithelial cells), the influence of periodontal treatment on the periopathogen loads is still not clear. Our preliminary data from RNAseq analysis identified Pg-(gingipain RgpA and RNA polymerase sub-unit beta) and Aa-encoded (Elongation Factor–Tu, Ribosome protein L2) to be the transcripts that were highly expressed in gingival samples obtained from periodontitis subjects. This expression was consistent across three different patient cohorts across distinct geographic locations. Hence, we assessed the changes in these bacterial transcripts after non-surgical periodontal therapy. Gingipains that are involved in predisposing attachment and colonization of *P. gingivalis* ^23,24,26^ and aiding its co-aggregation with other periodontal pathogens in the biofilm were not detected in the healthy cohort and after periodontal therapy. Likewise, rpoB that is responsible for the bacterial RNA synthesis was also not detected in healthy subjects and exhibited fairly undetermined levels after therapy. Genes encoded by Aa namely EF-Tu and RPL2 were detected in all three groups. However, their expression levels were minimal in healthy subjects and after non-surgical periodontal treatment. This observation could be owing to the fact that *A. actinomycetemcomitans*, an opportunist oral commensal member has been detected in at-least one-third of the healthy subjects ^25,23^. EF-Tu, one of the most abundant proteins in the bacterial cells and RPL2 which is the primary RNA binding protein; can confer antibiotic resistance and alter translation frequency, thus aiding in the defense of this organism by synergistic interactions in the biofilm and facilitate a protected survival in the gingival crevice ^23,27,28^. Thus, the reduction in bacterial loads by the way of reduced periopathogen encoded transcripts corroborated with clinical improved outcomes.

Periodontal therapy triggers periodontal tissue regeneration, requiring a change in the cellular milieu and a micro-environment supportive of resolution of inflammation. Our therapy clinically validated the assessment of M1/M2 polarization markers during the resolution of periodontal inflammation. The increased levels of M1 markers and pro-inflammatory cytokines are in concordance with other reports evaluating the role of MΦ polarization ^29–31^. Bonafide evidence of *in vivo* macrophage alternative activation, which elicits a strong Th2 response; was evidenced by increased IL10 and CCL22 expression, in a more comprehensive functional perspective after non-surgical periodontal therapy. CCL22, also known as macrophage-derived chemokine (MDC) is mainly produced by macrophages upon the stimulation with microbial products, is upregulated by Th2-type cytokines to enhance the chemotactic migration and recruitment of DCs and Th2 cells ^32^. In addition to the host immune responses in retaliation to the imbalances in periodontal microbiota; environmental and epigenetic factors for e.g. miRNAs are also responsible in directing the fate of macrophage M1, M2 switch in periodontal disease ^33–35^. In our study, we noted higher expression of M1 markers and lower M2 markers in inflamed gingiva, which improved remarkably post-therapy suggesting dynamic molecular changes in immune cells polarization and activity after a short 4-6 week period of treatment. Monitoring these molecular profiles can serve as a promising approach in periodontal treatment. For instance, M1 associated markers STAT1, miR-155 or the effector cytokines / chemokines TNF-α or CXCL10 that were responsive to periodontal therapy can be used to target specific subset of population with exaggerated immune responses. Conversely, M2 associated markers STAT6, TGM2 or the effector cytokines / chemokines IL10, CCL22 that were elevated after therapy, can be utilized in evaluating response to therapy. Using our murine LIP and resolution model, we further validated our human findings of perturbation in M1 and M2 phenotype in periodontal disease. Our results demonstrate that macrophages are highly response to periodontal immune microenvironment and swiftly shifts towards reparative M2 phenotype upon removal of dysbiotic trigger (ligature in this case). Early detection and intervention utilizing effective biomarkers can help minimize or prevent periodontal destruction. Decrease in the levels of M1Mφ markers and conversely, the increase in M2Mφ polarization markers correlate with clinical improvement in periodontal disease.

The tissue equilibrium achieved between the microbes in the plaque biofilm in periodontal health may easily disrupted in the states of chronic inflammatory conditions like periodontal disease and lead to host-immune mediated tissue damage. The deranged immunomodulatory responses could occur via over-activation of either pathways. Our results highlight the significant role for immune governed mechanisms by MΦ plasticity and polarization in oral inflammation. An innovative and noteworthy finding was the restoration of M2 marker expression levels post-therapy to similar levels observed in periodontally healthy subjects, as well as reduction in the M1 marker expression levels that corroborated with clinical improvement. This further affirmed the immune mediated restoration and/or maintenance of periodontal tissue homeostasis. It is the inherent plasticity that adeptly changes according to the tissue micro-environment that determines M1 to M2 switch and vice versa. This MΦ plasticity is essential to combat pathogen/antigen clearance as well as instigate tissue repair ^36,37^.

## Conclusion

This is one of the first translational studies to longitudinally assess the M/M2 MΦ profiles before and after therapy in periodontitis subjects and can serve as the first step in examining current treatment strategies to include investigations on the role of macrophage immune mediated regulatory networks governing periodontal disease pathogenesis. Our findings highlight the immuno-modulatory role of M1/M2 MΦ polarization in the periodontal disease pathogenesis and likewise, the impact of periodontal therapy in causing a switch towards M2 MΦ phenotype. Assessment of M2 macrophage markers may provide clinically relevant information to evaluate patient response to periodontal therapy and may be useful to target non-responders with exaggerated immune responses.

## Acknowledgements

This study was supported by the NIH/NIDCR R01 DE027980 and R03DE027147 to ARN.

